# FRAGMENTED BIODIVERSITY: FERNS AND LYCOPHYTES FROM FOREST FRAGMENTS IN JUIZ DE FORA, MINAS GERAIS, BRAZIL

**DOI:** 10.1101/2021.02.17.431701

**Authors:** Lucas Vieira Lima, Vinícius Antonio de Oliveira Dittrich, Filipe Soares de Souza, Cassiano Ribeiro da Fonseca, Alexandre Salino

## Abstract

The Atlantic Forest is one of the most threatened formations in the world. In this context, the urbanization process stands out as one of the major factors causing environmental degradation, mainly due to the loss of native vegetation and habitat destruction. In order to fill this gap, we carried out the inventory of ferns and lycophytes in the forest remnants of the municipality of Juiz de Fora. We analyzed more than 1,353 samplings recorded throughout approximately 150 years, as result we recorded a total of 174 species distributed in 73 genera and 26 families. The most representative families were Pteridaceae with 32 species, followed by Polypodiaceae with 26 and Thelypteridaceae with 20. In addition, we present the historical data on the sampling of ferns and lycophytes, as well as the history of the fragmentation process of the Atlantic Forest remnants in the municipality. Juiz de Fora becomes an interesting model for broader floristic studies, generating subsequent subsidies for conservation actions and preservation of the natural patrimony.

## RESUMO

A Floresta Atlântica é um dos domínios mais ameaçados do planeta. Neste contexto, o processo de urbanização configura-se como um dos maiores fatores causadores da degradação ambiental, fundamentalmente pela extinção da vegetação nativa e destruição de habitats. Visando contribuir para o preenchimento desta lacuna, foi realizado o inventário das samambaias e licófitas nos remanescentes florestais do município de Juiz de Fora. No total, foram analisados mais de 1353 registros de coletas realizadas no intervalo de aproximadamente 150 anos. Foram registradas 174 espécies, distribuídas em 73 gêneros e 26 famílias. As famílias mais representativas foram Pteridaceae com 32 espécies, seguida de Polypodiaceae com 26 e Thelypteridaceae com 20. Adicionalmente, são apresentados dados históricos de coletas de samambaias e licófitas, bem como o histórico da fragmentação dos remanescentes de Floresta Atlântica no município. Deste modo Juiz de Fora torna-se um interessante modelo para estudos florísticos mais amplos, gerando assim posteriores subsídios para ações de conservação e preservação do patrimônio natural da humanidade.

## INTRODUCTION

Ferns and lycophytes are two distinct lineages of seedless vascular plants, sharing a life cycle in which the gametophytic phase is independent from the sporophytic one (Page, 1979,; PPG I, 2016). In a recent review of the classification of ferns and lycophytes (PPG I, 2016), it has been estimated that there are 11,916 species. One of the main centers of diversity and endemism is the neotropical region (Moran 2008), with much of the species richness and endemism associated with the mountains of southeastern Brazil, Mexico, and the Andes (Tryon, 1972; Moran 2008).

Almeida & Salino (2016) demonstrated that, despite the current favorable scenario for the use of new molecular and computational techniques in systematic and biogeography studies, there is still a large gap in the knowledge about the diversity of groups of ferns and lycophytes. Among the factors related to the lack of knowledge about the diversity and patterns of geographic distribution of ferns and lycophytes, the main ones are the degradation, fragmentation and destruction of habitats in megadiverse neotropical countries such as Brazil (Almeida & Salino, 2016, Cincotta *et al*., 2000; Meyer *et al*., 2000).

The Atlantic Forest domain is an important set of ecosystems with high levels of endemism and richness (Myers *et al*., 2000, Stehmann *et al*., 2009), considered as one of the 34 global biodiversity hotspots (Mittermeier *et al*., 2004). Due to a long history of exploitation and degradation, currently only 7-8% of the original coverage remains (Galindo-Leal & Câmara, 2005). More recent studies using remote sensing methodologies with a resolution of 5 km^2^ demonstrated the recovery of vegetation cover in the Atlantic Forest, reaching 28% of the original coverage (Rezende *et al*., 2018). However, these remaining areas are constituted by mosaics of small and biologically impoverished fragments, on average smaller than 100 ha, whose restoration could take hundreds of years (Liebsch *et al*., 2008, Ribeiro *et al*., 2009, 2011).

The Seasonal Semideciduous Forest (SSF) is today restricted to a mosaic of forest fragments at different stages of regeneration (Whitmore, 1978, Ferreira Júnior *et al*., 2007), contrasting with its wide past distribution. According to Stehmann *et al*., (2009), this formation is the second most important in terms of plant richness and endemism in the Atlantic Forest domain. For fern and lycophyte species, 48% (406 spp.) of the total recorded for the Atlantic Forest (840 spp.) occur at SSF, of which about 33% (89) of them are endemic to the formation (Salino & Almeida, 2009a).

The municipality of Juiz de Fora, located in the southeastern portion of the state of Minas Gerais, is predominantly under the domain of the Semideciduous Seasonal Forest (Lima & Dittrich, 2016; PMJF, 2017). The native forest cover was almost completely destroyed, initially by opening up areas for agricultural activities, with emphasis on the coffee culture, later transformed into pasture areas, mainly of molasses grass (*Melinis minutiflora* P. Beauv.) and more recently palisade or signal grass (*Urochloa* spp.). The few remaining forest fragments are represented mainly by secondary formations in different succession stages (Almeida, 1996).

In addition, the constant anthropic pressures in response to the increased demands of the human population constitute an imminent threat to the maintenance of biodiversity (Almeida & Salino, 2016). In the last 20 years, the population of Juiz de Fora has grown by around 24%, reaching approximately 560 thousand inhabitants (IBGE 2016). The municipality currently has about 28 thousand hectares of native vegetation cover, which represent approximately 20% of its territory (Scolforo & Carvalho, 2006, Fonseca & Carvalho, 2012). However, only 4% of these forested areas are under protection in conservation units (Fonseca & Carvalho, 2012, PMJF, 2017).

The long history of exploitation and degradation to which the municipality’s vegetation is subjected and its importance in the representativeness of the Atlantic Forest fragments make it essential to carry out inventories of the flora of this area. In this context, the present study had as its main objective carrying out the inventory of ferns and lycophytes of the forest fragments in Juiz de Fora.

## MATERIALS AND METHODS

We reviewed all specimens of ferns and lycophytes from the studied area deposited in the BHCB and CESJ and herbaria (acronyms according to Thiers, 2020). In addition, we consulted and reviewed the identifications of specimens with images available in the following virtual herbaria: INCT-Herbário Virtual da Flora e dos Fungos (INCT-Virtual Herbarium of Flora and Fungi 2018), Jabot – Banco de dados da Flora Brasileira (JBRJ, 2017), and Reflora -Virtual Herbarium (Reflora, 2017). In a total we reviewed 1353 records of ferns and lycophytes from all forest remnants of Juiz de Fora.

The municipality of Juiz de Fora is located in the southeast of the state of Minas Gerais, between the approximate geographical coordinates of 21° 31’ 16” and 21° 59’ 59” S and 43° 08’ 50” and 43° 41’ 10” W. The regional climate can be characterized as Köppen type Cwa (mesothermal, with hot and rainy summers). The average annual precipitation, measured between the years 1999 and 2011, is 1627.8mm and the average annual temperature is around 19.3 ° C, for the same period. The altitude varies from 467 to 1104m and the relief is diverse, with concave-convex hills and valleys (Anuário Estatístico de Juiz de Fora, 2012). The municipality has remaining areas of Atlantic Forest, permeated by the city. Around 24% (9,662 ha) of the urban area of the municipality of Juiz de Fora is covered by forests, distributed in 1,122 fragments. Consequently, the average size of these fragments is 8.61 ha (Barros 2015). Most of these fragments have different sizes, are at different stages of regeneration and still suffer constant impact from anthropic action.

We used ArcGIS ver. 10 (ESRI 2011) to build a map showing the forest areas of Juiz de Fora (Figure 2). We adopted PPG I (2016) as the classification system, and follow the definitions of Lellinger (2002) and Zuquim *et al*., (2008) to categorized the species’ preferential substrate in: terrestrial, rupicolous, corticicolous, hemicorticicolous, amphibians, and aquatic.

**Figure 1.**
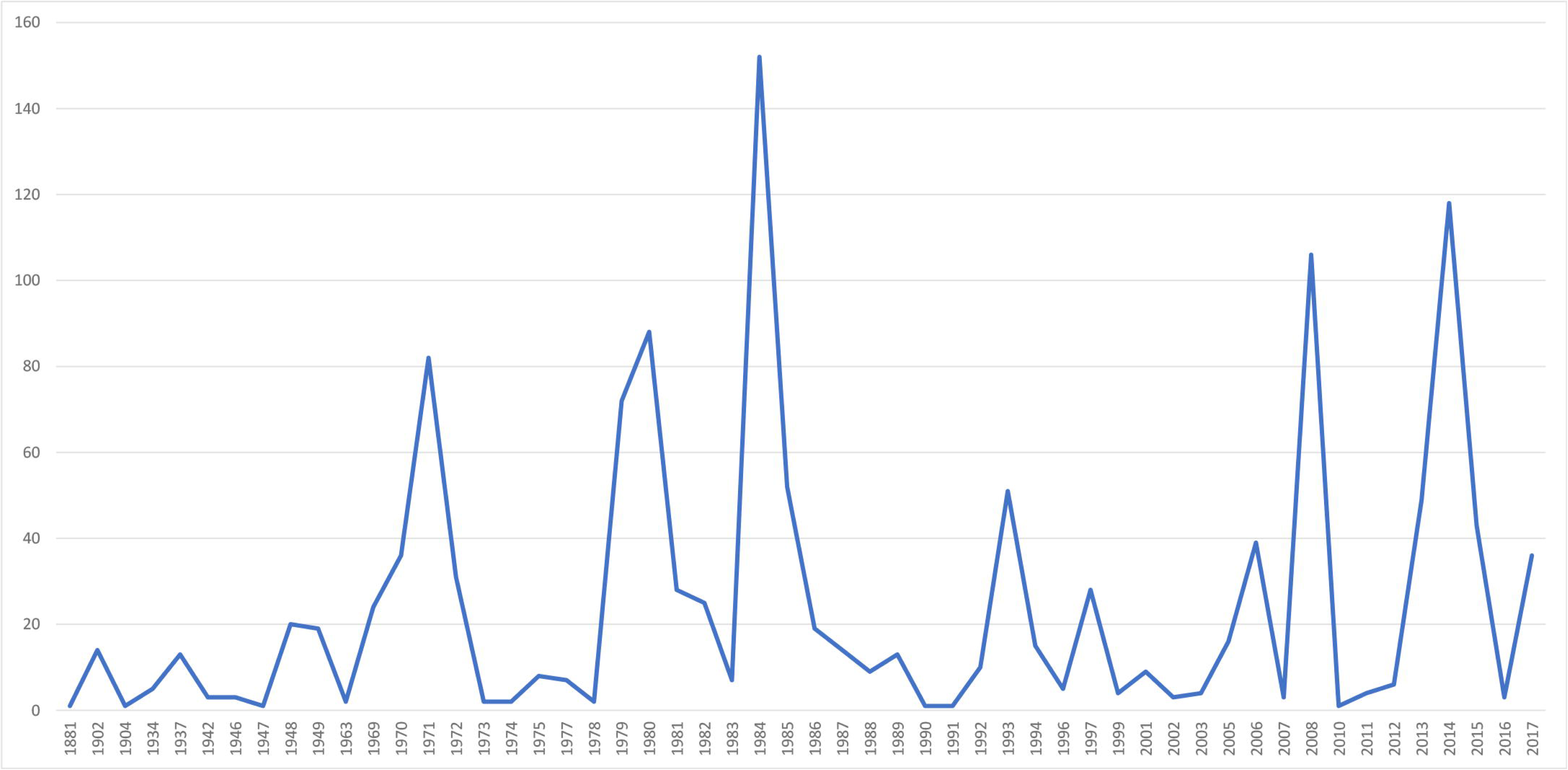
Number of samples of ferns and lycophytes in Juiz de Fora though the years.

**Figure 2.**
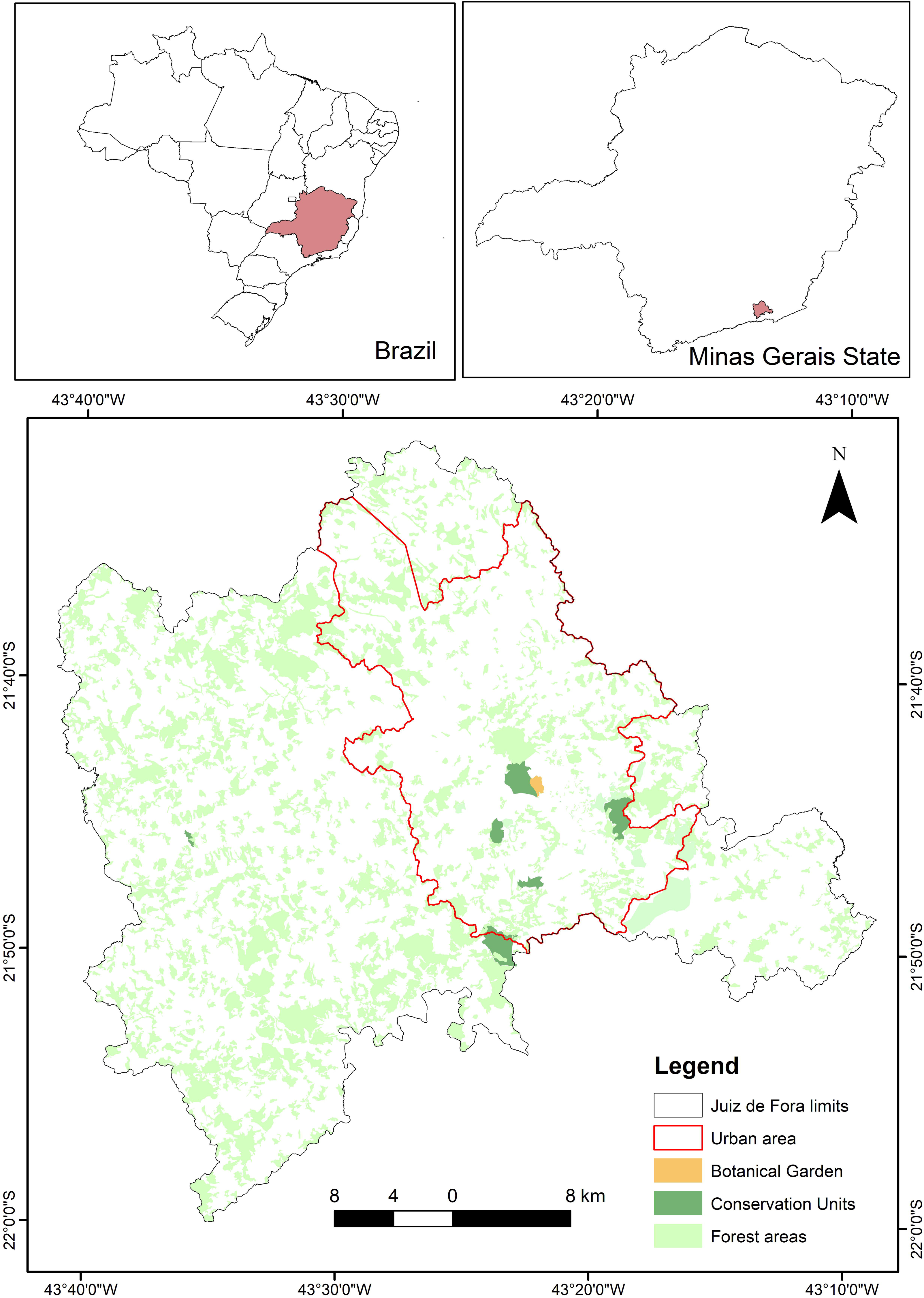
Map showing the forest areas of Juiz de Fora.

## RESULTS

### 1. Sampling history

The first samplings of ferns and lycophytes in the municipality of Juiz de Fora date from the second half of the 19th century. *Elaphoglossum nigrescens* was the first species to be sampled. Collected in February 1862, it was deposited at Herbário Capanema in Manaus (Amazonas State). Currently the specimen is stored in the herbarium of the Botanical Garden of Rio de Janeiro (RB) (RB00687659), and the voucher has an original label dated 1862, in addition to another label transcribed with the spelling of Alex Curt Brade (1881-1971), with the date changed to 1863.

At the beginning of the 20th century, other samplings were made by the Carl August Wilhelm Schwacke (1848-1904). The specimens came from one of Schwacke’s expeditions to the state of Minas Gerais, passing through Juiz de Fora between June and July of 1902. The locations sampled by Schwacke in Juiz de Fora were Poço d’Anta and Morro do Imperador, where 14 species were sampled, most of which are stored at the RB herbarium. Subsequently, the ferns and lycophytes of Juiz de Fora was explored by Brade in two sampling expeditions, in 1934 and 1937 respectively, in which another 14 species were recorded for the municipality.

From the first sampling to the first half of the 20th century, although the municipality of Juiz de Fora had great importance in the national context for coffee production and its industries, the knowledge local flora remained incipient. The second half of the 20th century was marked by a significant increase in samplings, driven by the founding of the Leopoldo Krieger Herbarium (CESJ) in 1940, as an initiative of the priests Leopoldo Krieger (1919-2008) and Luiz Roth, by the time students of Theology (Salimena & Menini-Neto, 2008).

The first samplings by Krieger and Roth in Juiz de Fora date from the years 1942, 1948, and 1949. These are single samplings of which only 28 specimens were incorporated into the newly founded herbarium. In 1969 Krieger was hired as a professor at the Universidade Federal de Juiz de Fora (Salimena & Menini-Neto, 2008). From this date, a significant volume of samplings was recorded for the municipality, done by the priest and his students until the end of the 1980s. Of the 1,353 sampling records for the municipality, approximately 672 (49%) were collected by Krieger, who was the first sampler in more than 90% of the cases, between 1969-1989. From 1990 to the present, with the Biology course at Universidade Federal de Juiz de Fora already strongly consolidated, samplings started to be done by a more varied number of samplers and the number of records practically doubled.

### 2. Floristic composition

We found 175 species, distributed in 73 genera and 26 families. The richest families were Pteridaceae with 32 species and eight genera (18% of species), Polypodiaceae with 26 species and eight genera (14%) and Thelypteridaceae with 20 species and seven genera (11%) (Figure 1, Table 1). The most representative genera were *Adiantum*, with nine species, *Amauropelta, Asplenium, Microgramma*, and *Pteris*, with seven each, *Cyathea* and *Selaginella*, with six each, *Anemia, Ctenitis, Doryopteris, Pleopeltis*, and *Serpocaulon*, with five species each. Regarding the habit, about 16% of the species are epiphytes and about 84% are terrestrial or rupicolous. In addition, naturalized species such as *Deparia petersenii, Macrothelypteris torresiana, Christella dentata*, and *Pteris vittata* were collected.

**Table 1.**
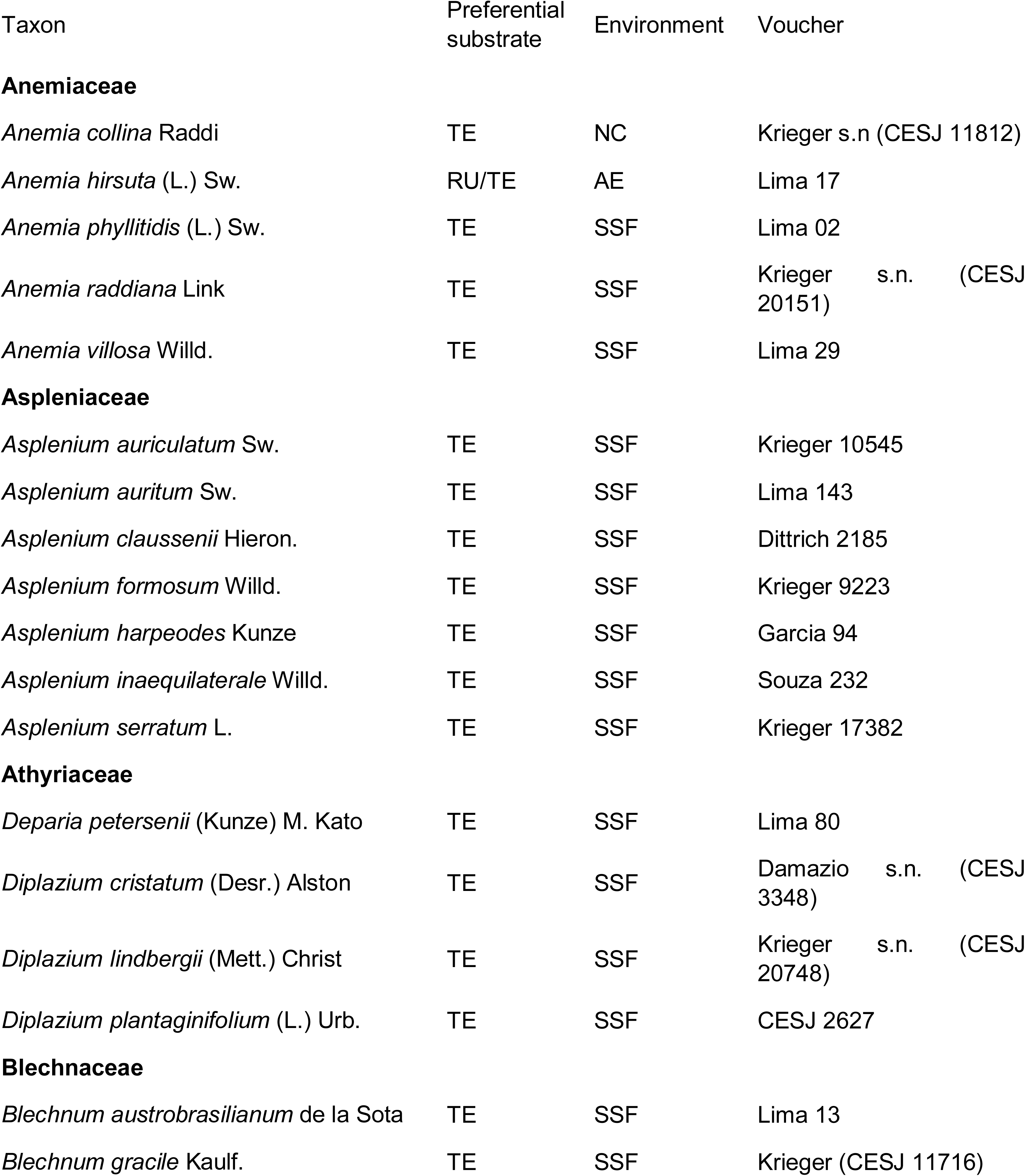
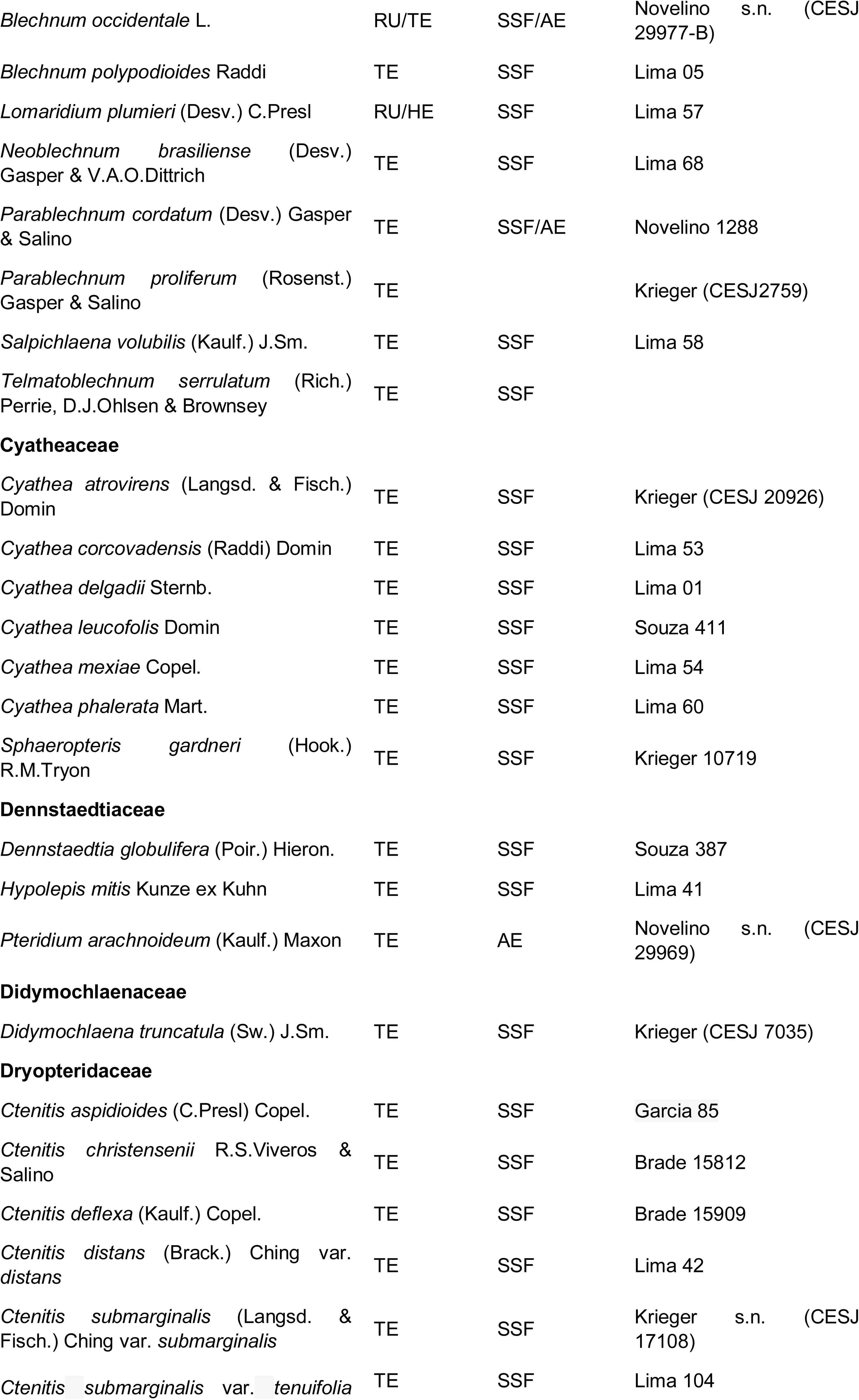
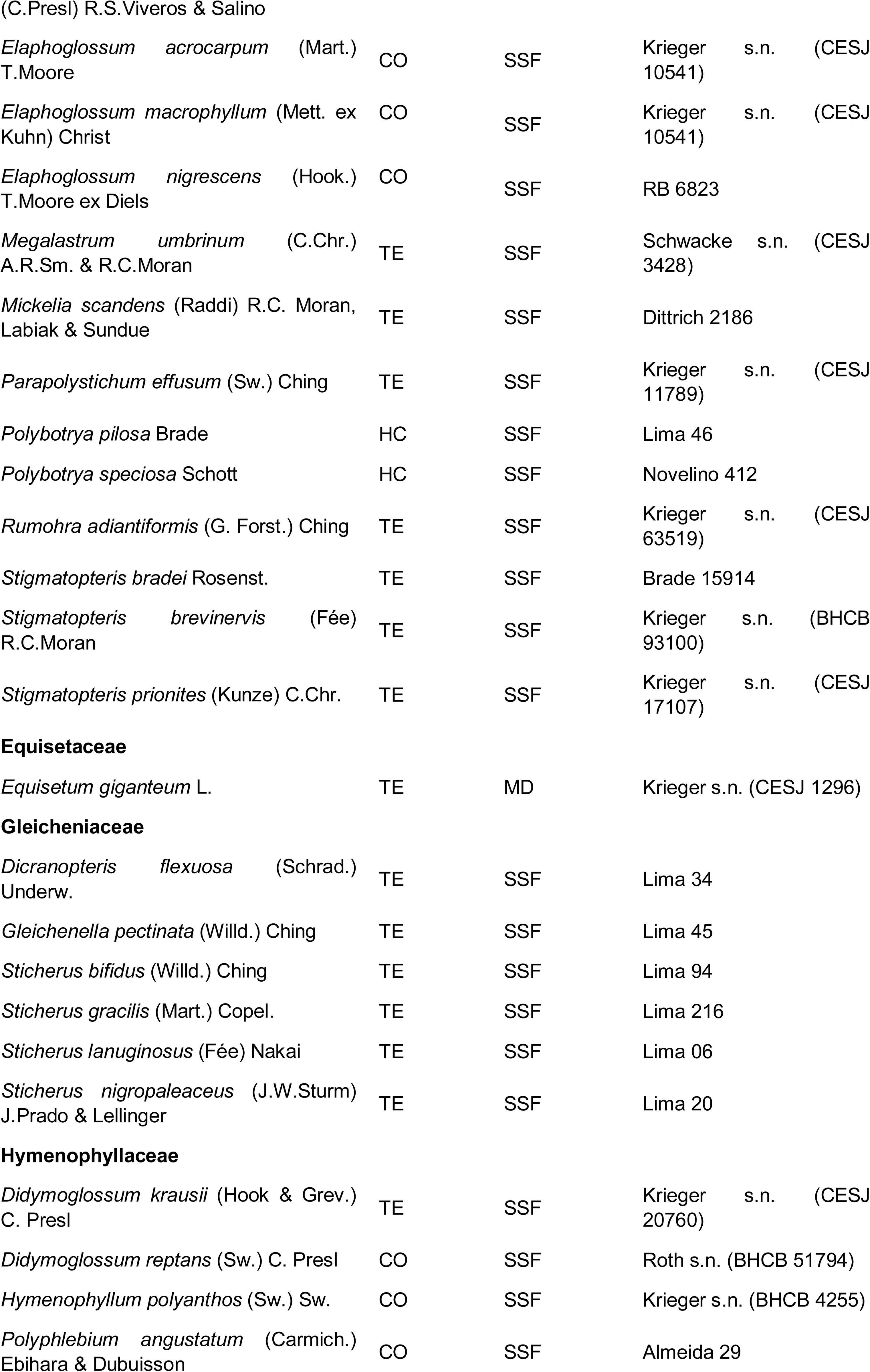
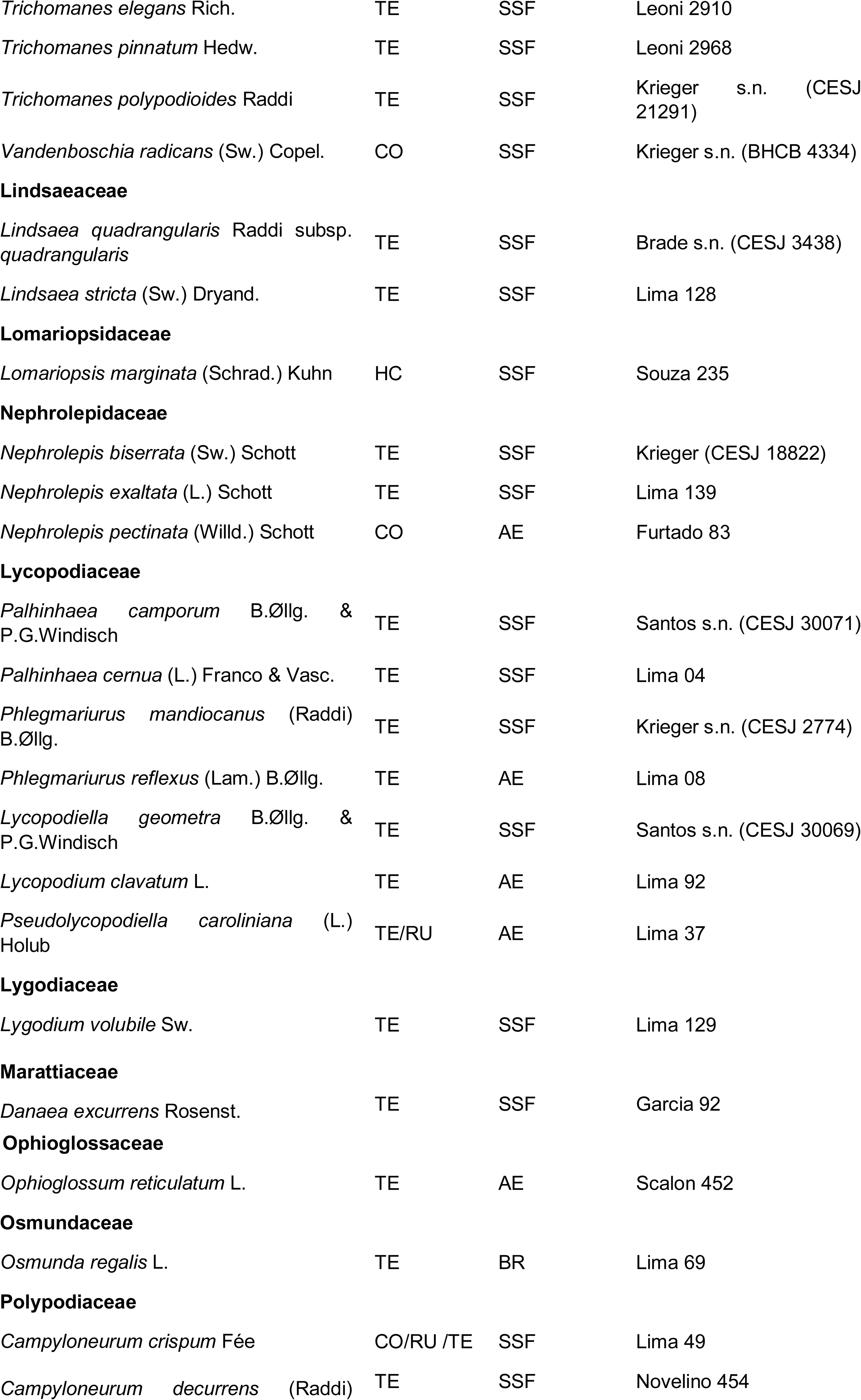
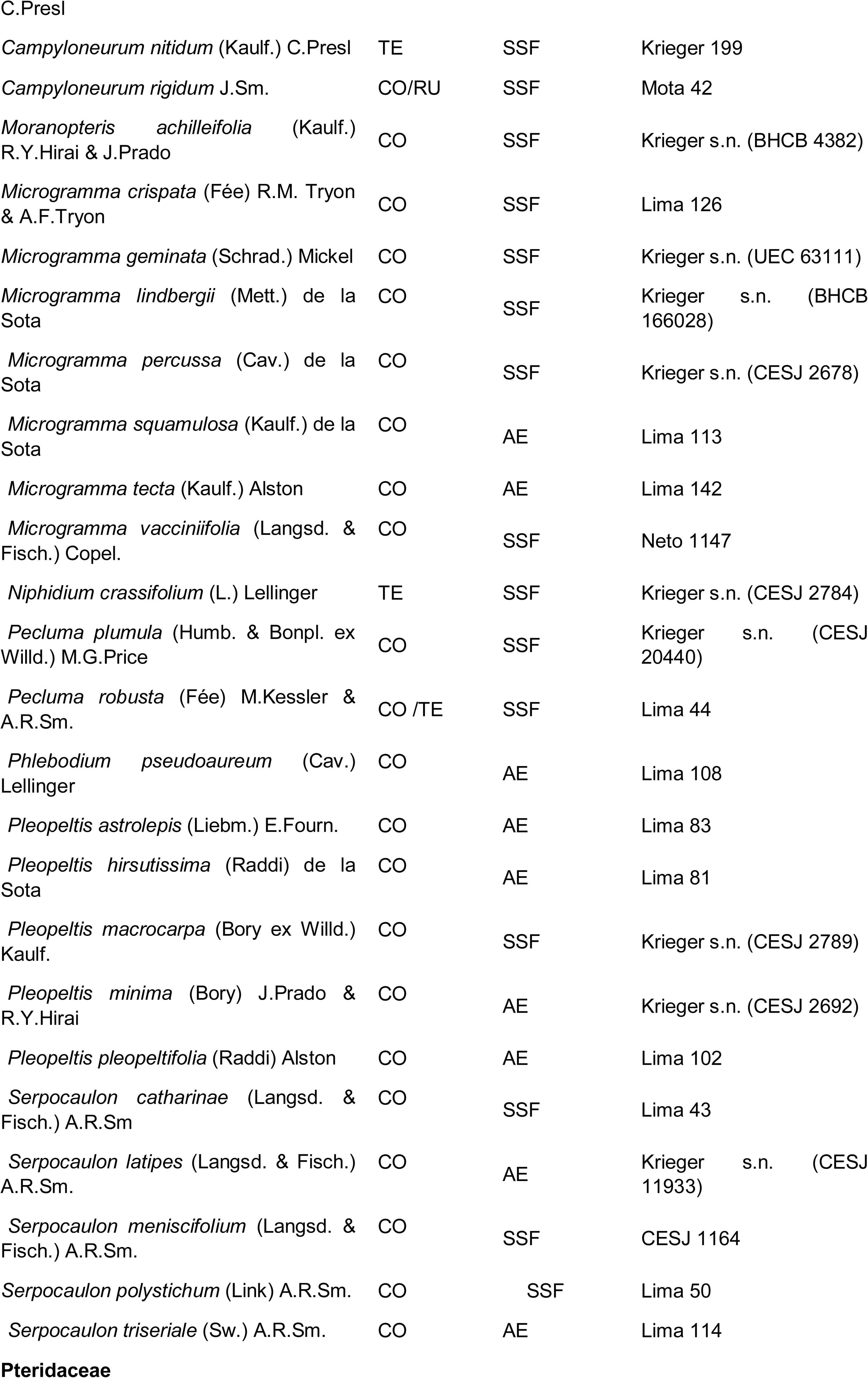
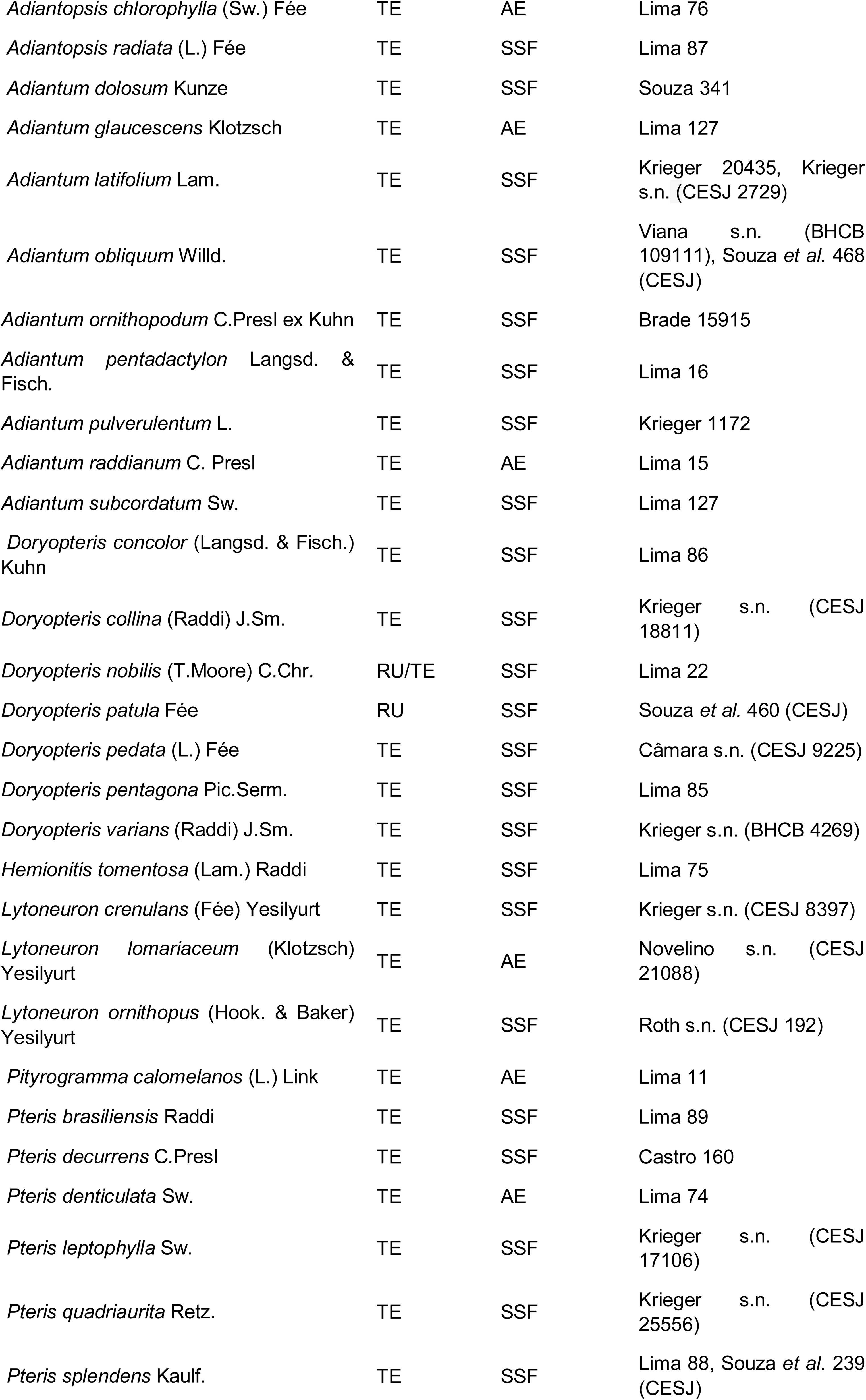
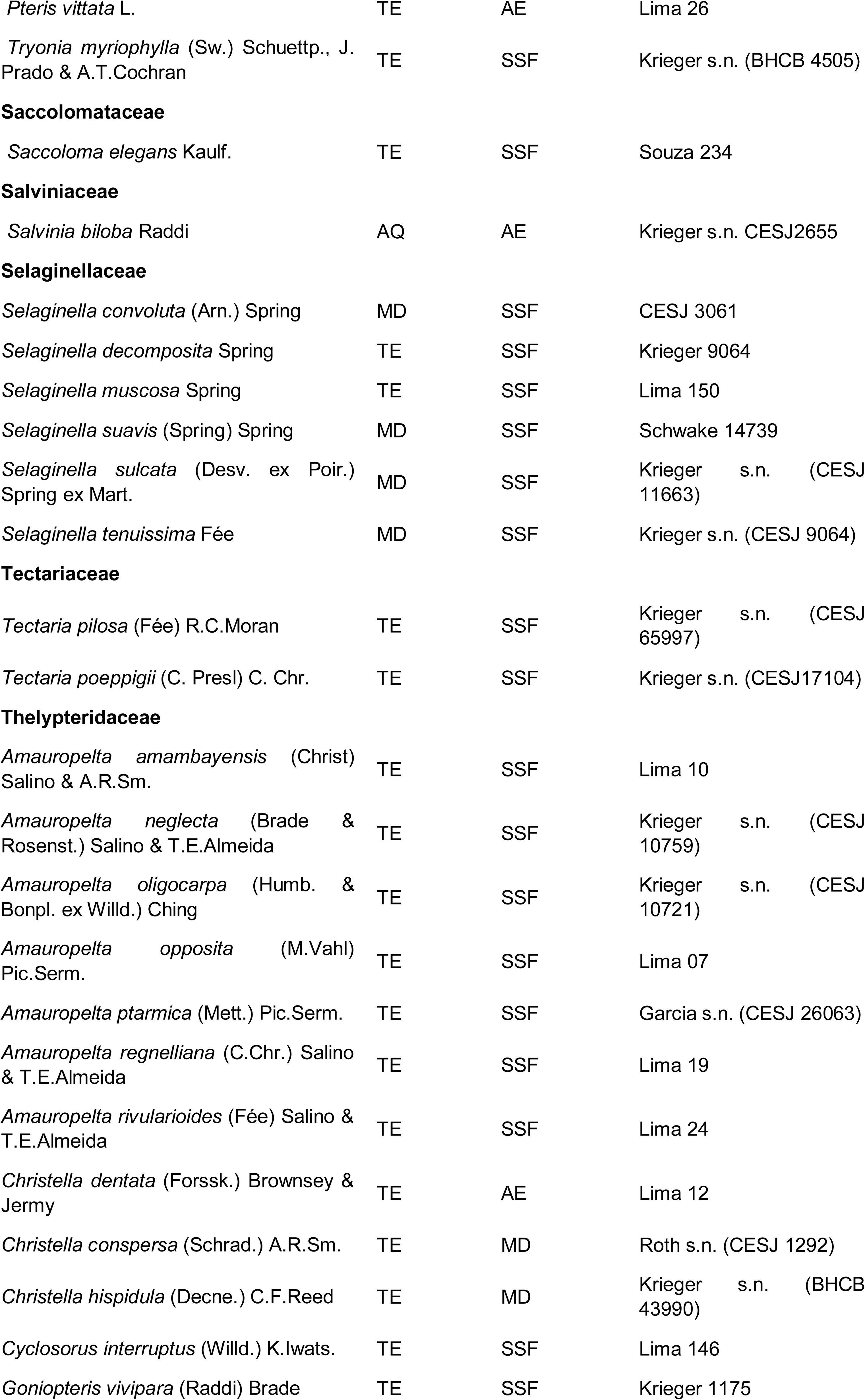
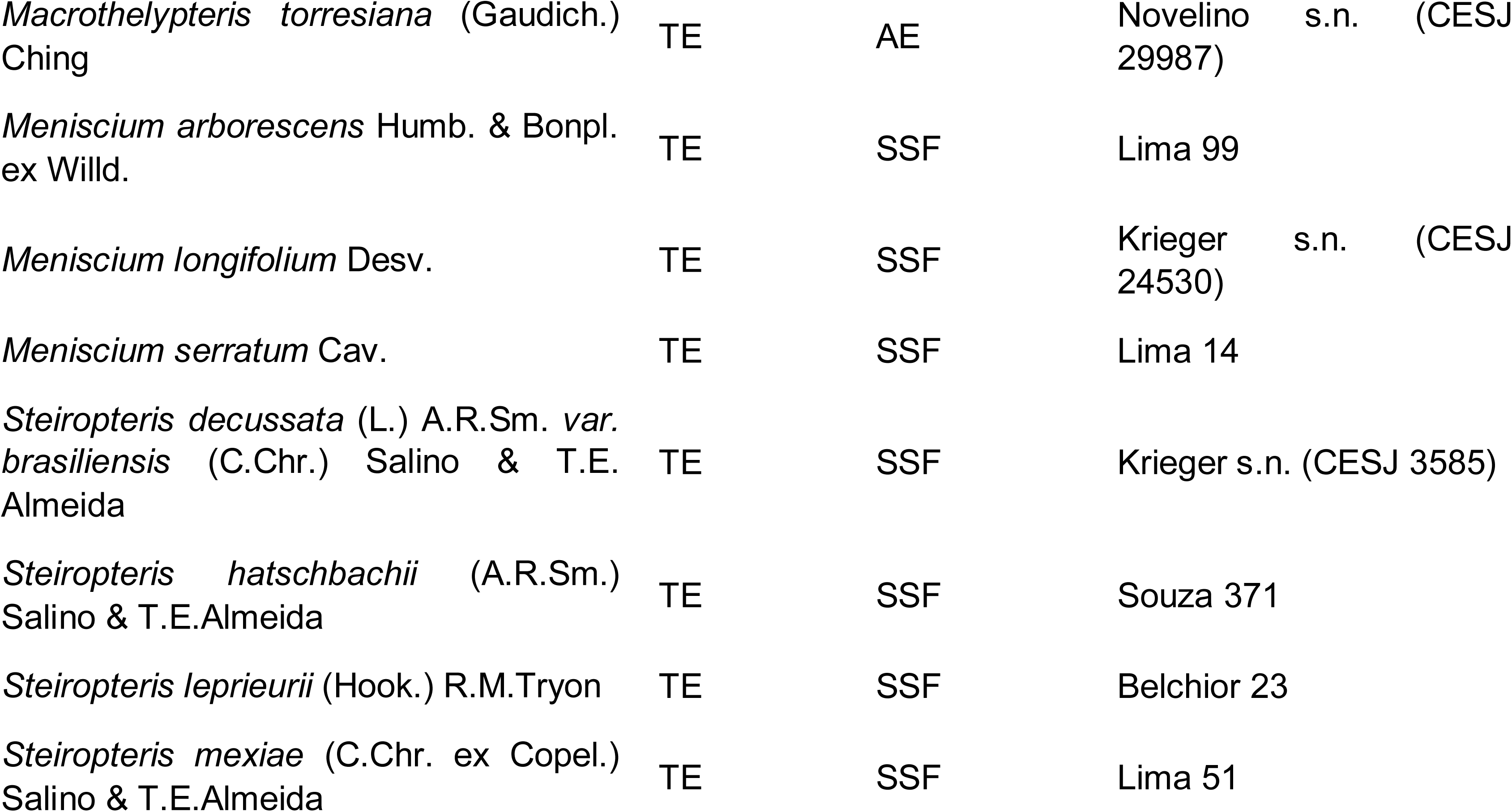
Ferns and lycophytes species in the municipality of Juiz de Fora. AE: anthropogenic environment, CO: Corticicolous, HC: Hemicorticicolous, MD: missing data, NC: not computed, RU: rupicolous, SSF: Seasonal Semideciduous Forest, TE: terricolous.

## DISCUSSION

Salino & Almeida (2009a) pointed out, among the main threats to species of ferns and lycophytes in the State of Minas Gerais, the decrease and degradation of habitats, due to the subsequent decline in the optimum conditions for establishment and survival. Nevertheless, Lima & Dittrich (2016) recorded 78 species in only three remaining areas of Atlantic Forest in Juiz de Fora, which represent nearly 43% of the total registered here. This shows that few areas, even if immersed in an urban matrix with more than 500 thousand inhabitants, may house a high number of species of native flora.

The results of the present study point to a higher number of species compared to inventories carried out in other areas of the Southeast and South of Brazil in which the main phytophysionomy is composed of Seasonal Semideciduous Forest (SSF) (Table 2). Colli *et al*. (2004b) recorded 34 species for Seasonal Semideciduous Forest from Parque Estadual de Vassununga, São Paulo State, in an area of 1,732 ha with an altitude between 500-750m. Melo & Salino (2002a) recorded 116 species for Parque Estadual do Rio Doce (PERD), which has a larger area with 35,970 ha. Despite the large area of forest of PERD, less fragmented than the forest formations of Juiz de Fora, the low number of species found may be related to the low altitude and limited altitudinal variation of PERD (215 to 525m). Salino & Almeida (2009a) showed that altitudinal ranges above 700 m are the richest in the state of Minas Gerais. Similarly, Senna & Kazmirczak (1997) recorded 45 species in an area of 1,031 ha of Seasonal Semideciduous Forest in Morro da Extrema, Rio Grande do Sul State. In this case, the low number of species may be associated with the maximum altitude of the area at 225m (Table 2). Furthermore, the southern region of Brazil has fewer species of ferns and lycophytes than the southeastern region (Flora do Brasil 2020 Online 2020).

**Table 2.** Comparison among studies that inventoried the Seasonal Semideciduous Forest (SSF) areas in the Atlantic Forest.

| Reference | Locality | Vegetation formations | Area (ha) | Elevation (m) | Total of spp. |
| --- | --- | --- | --- | --- | --- |
| Present study | Juiz de Fora | SSF | 142988 | 467-1104 | 181 |
| Salino *et al.* 2013 | Parque Estadual do Rio Preto | SSF and Cerrado | 12000 | 750-1850 | 185 |
| Colli *et al.* 2007 | Reserva Estadual de Águas da Prata | Riparian Forest and SSF | 48,40 | 840-1060 | 39 |
| Colli *et al.* 2004a | Floresta Estadual de Bebedouro | Riparian Forest, Cerrado, and SSF | 99,5 | 570 | 10 |
| Colli *et al.* 2004b | Parque Estadual da Vassununga | SSF | 1732 | 500-750 | 34 |
| Souza *et al.* 2012 | Serra Negra | Rain Forest, Highland Fields, and SSF | 10000 | 900-1698 | 209 |
| Figueiredo & Salino 2005 | Patrimônio Natural ao Sul da Região Metropolitana de Belo Horizonte | Cerrado, Ferruginous Fields, and SSF | 2280 | 850-1420 | 190 |
| Melo & Salino 2002 | Estação Biológica de Caratinga | SSF | 880 | 400-680 | 102 |
| Melo & Salino 2002 | Parque Estadual do Rio Doce | SSF | 35970 | 230-515 | 116 |
| Melo & Salino 2007 | APA Fernão Dias | Rain Forest and SSF | 180 | 1000-2068 | 173 |
| Nobrega & Prado 2008 | Jardim Botânico Municipal de Bauru | Cerrado, Riparian Forest, and SSF | 321,71 | 510-540 | 54 |
| Senna & Kazmirczak 1997 | Morro da Extrema. Porto Alegre | SSF | 1031 | Up to 255 | 45 |

Melo & Salino (2002b) recorded 102 species in an area of 880 ha of SSF at Estação Biológica de Caratinga, Minas Gerais. This area has medium altitudes, and the forest remnants are more continuous than those of Juiz de Fora (JF). It is to be expected that in inventories on a regional scale, as performed in JF, the number of species will be greater than inventories on a local scale (Mehltreter 2010). However, the lower number of species found when we observe the species/area ratio may be associated with the broad history of exploitation, degradation, and fragmentation of forest formations in Juiz de Fora.

Similarly, Figueiredo & Salino (2005) recorded 190 species in fragmented areas in the metropolitan region of Belo Horizonte, Minas Gerais. Such areas comprise a mosaic of phytophysiognomies in a transition zone between Cerrado and Atlantic Forest, of which the predominant phytophysionomy is Seasonal Semideciduous Forest. This work demonstrates that, as in Juiz de Fora, such fragmented areas, although under constant threat, can be important refuges and harbor a considerable number of native species.

According to Salino & Almeida (2009a), in their diagnosis of the ferns and lycophytes flora of the Minas Gerais State, the SSF was the forest formation with the highest number of species (429 spp. - 62.5%), exceeding the number of species recorded for the Tropical Rain Forest (384 spp. - 56%). The authors also correlate these factors to the relief of the state, since in Minas Gerais the domain of the Semideciduous Forest is mainly restricted to mountainous regions, where there are many deep valleys that allow the occurrence of most ferns and lycophytes.

Here 16 species for SSF listed by Salino & Almeida (2009b) as exclusive of the Atlantic Tropical Rain Forest (Table 1). These new records may be associated with some factors related to sampling deficiency and the fluctuation of micro-habitats due to variations in the relief forms of the studied area. Examples of these species are representatives of Hymenophyllaceae, registered mainly in the Rio do Peixe region, and *Danaea excurrens* (Marattiaceae), found in a valley bottom. On the other hand, it is interesting to note that Pteridaceae is the richest family (33 spp., Table 1), surpassing Polypodiaceae, Dryopteridaceae and Thelypteridaceae, a pattern that traditionally occurs in SSF formations (Melo & Salino, 2002; Salino & Almeida, 2009).

Regarding the rates of epiphytes, the present study found 16%, which seems to be close to the rates found in other SSF inventories such as Colli *et al*. (2007), for Reserva Estadual de Águas da Prata (15%), Colli *et al*. (2004b), for Parque Estadual de Vassununga (20%), and higher than the values found by Melo & Salino (2002a) for Estação Biológica de Caratinga (8.5%), Melo & Salino (2002b) for Parque Estadual do Rio Doce (8.2%), and Senna & Kazmirczak (1997) for Morro da Extrema (8%). These indexes are in accordance with the pattern found for this type of forest formation (SSF), with the regeneration stages of forest fragments, and may be associated with seasonality influencing the air humidity indexes, which is essential for the establishment of epiphytic plants (Figueiredo & Salino 2005).

Among the native species recorded, *Elaphoglossum acrocarpum* (Dryopteridaceae) stands out as threatened with extinction in the vulnerable category according to Kieling-Rubio *et al*. (2013). However, a few species of ferns and lycophytes can favor themselves in anthropized environments and increase the abundance of their populations, such as *Lycopodium clavatum* and species of Gleicheniaceae, such as *Dicranopteris flexuosa* and *Gleichenella pectinata*. Such plants occupy anthropized and altered areas like ravines by the roadsides. In addition, naturalized species can also benefit from environmental change and be widely spread, such as *Macrothelypteris torresiana, Christella dentata, Pteris vittata*, and *Deparia petersenii*, all found in Juiz de Fora. Finally, species of Gleicheniaceae and *Pteridium* (Dennstaedtiaceae), like the ones record to JF, can directly influence the process of succession and regeneration of secondary forests, both delaying the process and competing directly with native species (Mehltreter 2010).

Habitat destruction has a direct impact on the diversity of most species of lycophytes and ferns and may even lead to the extinction of local populations (Salino & Almeida 2009a, Mehltreter 2010). Through the observation of the extensive sampling history of more than 150 years, it is possible to assess that the frequency of sampling of some taxa changes. The three species of the genus *Elaphoglossum* (*E. acrocarpon, E. nigrescens*, and *E. macrophyllum*) recorded here were sampled only once, which may signal the imminent risk to the conservation of the local vascular flora.

Recently, a new species of Dryopteridaceae, *Ctenitis christensenii* R.S.Viveros & Salino, was described for the Brazilian Atlantic Forest and is here recorded for the first time for Juiz de Fora. The species was sampled only once in the municipality by Brade in 1937. Other than that, there is no other record of this species in the region, which means there is a great chance that this species may be locally extinct, as well as some other species. On the other hand, some species have a low frequency of collection due to specific characteristics of their biology such as *Ophioglossum reticulatum* - which is an annual plant - and *Pleopeltis minima*, which in addition to the small size presents poikilohydry, which makes it difficult to be spotted during the dry season.

Regenerating forest ecosystems hold only part of the original biodiversity (Paciência 2001), which is valid for forest fragments in the municipality of Juiz de Fora, which, despite the history of devastation and the different stages of regeneration, presents a high richness of ferns and lycophytes. The loss of biodiversity of the ferns and lycophytes in the studied areas may be associated with the deterioration of the optimum conditions of the environments, such as the changes in humidity and shading when compared to those offered in primary forests.

The regeneration of the studied forest fragments in the municipality of Juiz de Fora attests to the high resilience of the Atlantic Forest (Pinto & Brito 2003, Rezende *et al*. 2018), as exemplified by the regeneration of the Krambeck forest, which after being destroyed to form pastures, after 70 years of regeneration has shown significant recovery (Lima & Dittrich 2016). With the conservation of these forest areas over the years (progress in the succession process), the colonization of new species of ferns and lycophytes may occur and the abundance of more demanding species in terms of shade and humidity may increase (Figueiredo & Salino 2005). Therefore, the present results reaffirm the importance of maintaining and conserving urban forest fragments for sustaining Brazilian biodiversity in one of the biomes most affected by anthropic action.

## ACKNOWLEDGEMENTS

We thank Thaís Elias Almeida for notes and comments about this manuscript. This study was financed in part by the Coordenação de Aperfeiçoamento de Pessoal de Nível Superior - Brasil (CAPES) - Finance Code 001 (88887.19244/2018-00). We also thank CNPq for the research grant and scholarship (307115/2017-8) to A. Salino.

## Notes

### Competing Interest Statement

The authors have declared no competing interest.

